# Activation of the chemokine receptor CCR1 and preferential recruitment of Gαi suppress RSV replication: implications for developing novel RSV treatment strategies

**DOI:** 10.1101/2022.03.18.484968

**Authors:** Jiao Li, Ling Xue, Jiachao Wang, Aihong Meng, Jiajun Qiao, Miao Li, Xiuli Wang, Lingtong Meng, Jingyuan Ning, Xue Gao, Wenjian Li, Cuiqing Ma, Lin Wei

## Abstract

Respiratory syncytial virus (RSV) is a major pathogen that can cause acute respiratory infectious diseases of the upper and lower respiratory tract, especially in children, elderly individuals and immunocompromised people. Generally, following viral infection, respiratory epithelial cells secrete cytokines and chemokines to recruit immune cells and initiate innate and/or adaptive immune responses. However, whether chemokines affect viral replication in non-immune cells is rarely clear. In this study, we detect that chemokine CCL5 was highly expressed, while expression of its receptor, CCR1, was downregulated in respiratory epithelial cells after RSV infection. When we overexpressed CCR1 on respiratory epithelial cells in vivo/in vitro, viral load was significantly suppressed, which can be restored by the neutralizing antibody for CCR1. Interestingly, the antiviral effect of CCR1 was not related to type I interferon (IFN-I), apoptosis induction or viral adhesion or entry inhibition; in contrast, it was related to the preferential recruitment and activation of adaptor Gαi, which promoted inositol 1,4,5-triphosphate receptor type 3 (ITPR3) expression, leading to inhibited STAT3 phosphorylation, explicitly, phosphorylated (p)-STAT3 was verified to be among the important factors regulating the activity of HSP90, which has been previously reported to be a chaperone of RSV RNA polymerase. In summary, we are the first to reveal that CCR1 on the surface of non-immune cells regulates RSV replication through a previously unknown mechanism that does not involve IFN-I induction.

## INTRODUCTION

Respiratory syncytial virus (RSV) is the main cause of lower respiratory tract infection (LRTI) and infant respiratory diseases(1,2). RSV infection can lead to bronchiolitis and pneumonia and is associated with the development of recurrent wheezing and asthma(3). It has been estimated that 3.4 million people are hospitalized for RSV infection per year worldwide and that from 66,000 to 239,000 children younger than 5 years old die of LRTI caused by RSV. Among viral infections, RSV is considered to be the main pathogen threatening the health of elderly people(4-6).

The key factors determining the outcome of RSV infection are not fully understood, but the competition between the virus and host for survival is the crucial factor affecting the prognosis of RSV infection. Following viral infection, host cells recruit and activate immune cells by secreting a series of proinflammatory factors and chemokines to evoke innate and/or adaptive immune responses. Chemokines constitute a group of small-molecule proteins that are composed of 70 to 100 amino acids(7) and can promote the directional movement of immune cells. Despite the nonspecific function of chemokines, patterns of chemokine expression have been associated with specific disease states. In the field of respiratory viral infection, evidence has suggested that the response to viral invasion is regulated by a distinct chemokine expression profile that involves more CC chemokines than CXC chemokines(8).

Chemokines mediate a variety of biological effects by interacting with corresponding receptors on the cell surface. Most studies have mainly focused on the interaction between chemokines and receptors on immune cells, and immune cells are recruited to the infection site where they initiate an immune response. Notably, chemokine receptors are highly expressed in respiratory epithelial cells, begging the questions, what is the role of chemokine binding to host cell surface receptors, and does this binding have an impact on the prognosis of people with a viral infection? This study shows that when respiratory epithelial cells were infected with RSV, the expression of CCL5 increased significantly, while the expression of its receptor, CCR1, was downregulated. Moreover, the RSV load was increased continuously. Obviously, the downregulation of CCR1 expression inevitably weakens the ability of CCL5 to interact with host cells, which led us to wonder whether the interaction between CCR1 and CCL5 is related to RSV load. Therefore, we infected respiratory epithelial cells overexpressing CCR1 with RSV in vivo and in vitro. Interestingly, we found that with an increase in CCR1 overexpression, the viral load continued to decrease significantly compared with that in the control group. Ultimately, our study reveals a novel mechanism affecting the viral load in host cells; that is, the interaction between CCR1 and CCL5 affects the replication of RSV by regulating the activity of the RSV polymerase chaperone HSP90. This discovery may provide a new perspective on developing broad-spectrum antiviral drugs.

## RESULTS

### Following RSV infection, the expression of the chemokine CCL5 increased significantly and was negatively correlated with the replication of intracellular virus

In the study of RSV pathogenic mechanisms, Hep2 laryngeal cancer epithelial cells, A549 lung adenocarcinoma cells, BEAS-2B bronchial epithelial cells and Vero African green monkey kidney cells are the typically used models(9). In this study, 241 differentially expressed genes were identified by transcriptome sequencing analysis of RSV-infected A549 cells. Compared with that in the control group, the expression of 171 genes was upregulated and that of 70 genes was downregulated in the infected group (Fig. 1A). Among those thus identified, two genes in the CC chemokine family, CCL2 and CCL5, were very highly expressed (Fig. 1B), with CCL5 the most differentially expressed gene. Real-time quantitative PCR (qRT–PCR) verified that the chemokine CCL2, CCL3, and CCL5 expression levels gradually increased with prolonged viral infection, with the increase in CCL5 expression the most obvious (Fig. 1C). After RSV infection of three susceptible cell lines, A549, Hep2 and Vero cells lines, at a multiplicity of infection (MOI) of 0.5, the mRNA and protein expression levels of the chemokine CCL5 were significantly higher than those of the control groups within 48 hours (Fig. 1D and E). Additionally, 98 sputum samples were collected (RSV^-^, n=66, and RSV^+^, n=32) to detect the CCL5 mRNA expression level, and the results showed that the CCL5 mRNA level was significantly increased in the samples from patients with clinical RSV infection compared with that in the negative control sample (Fig. 1F). These results were consistent with the results of a gene chip analysis.

**Fig. 1.**
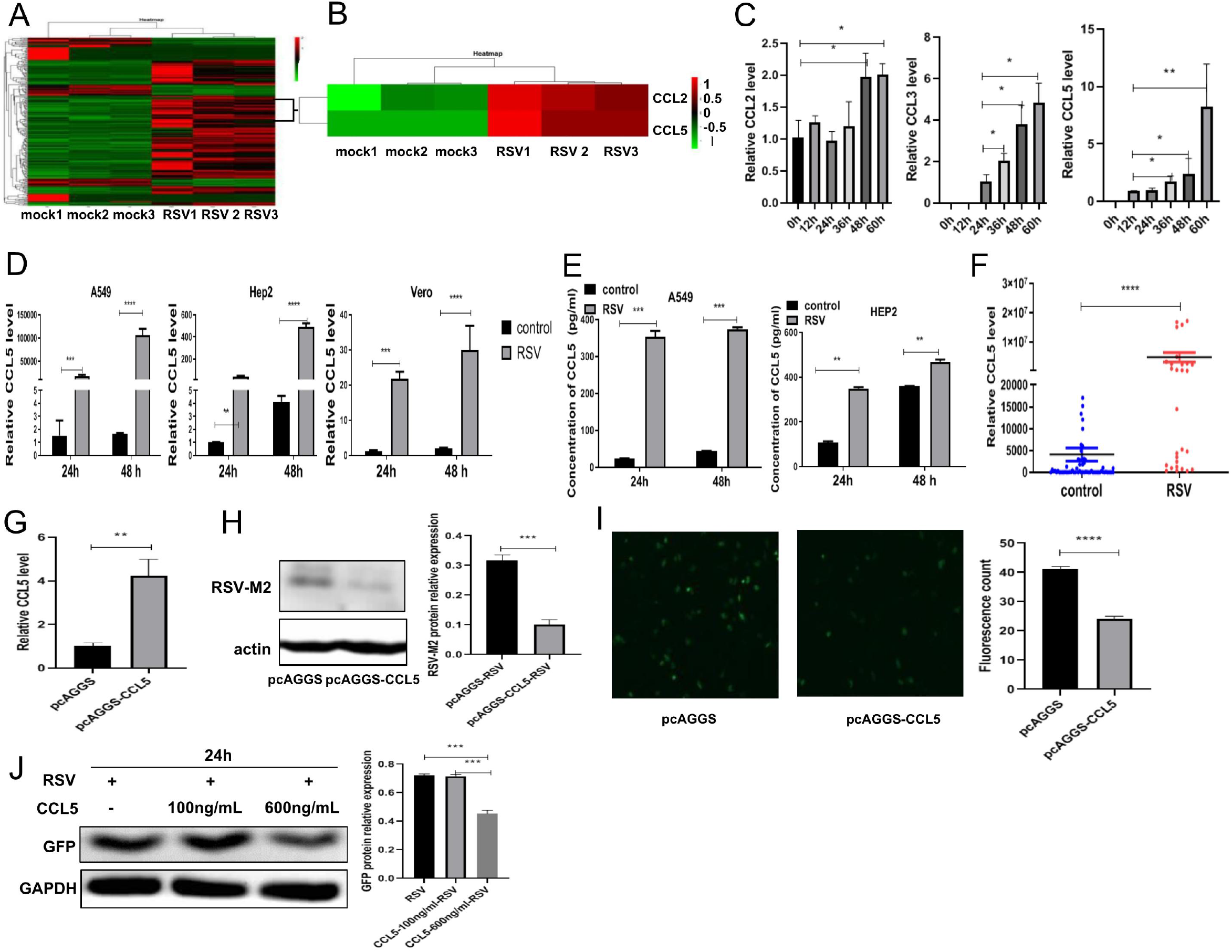
The expression of the chemokine CCL5 induced by RSV was increased. (A, B) Transcriptome sequencing was performed to analyze the expression of cytokines in A549 cells stimulated with RSV. (C) The expression of chemokines in A549 cells at different time points after RSV infection was analyzed by qRT–PCR. (D) The relative expression level of CCL5 mRNA in A549 cells, Hep2 cells and Vero cells infected with RSV was detected by qRT–PCR. (E) The expression level of CCL5 in the culture supernatant of A549 cells or Hep2 cells induced by RSV was measured by ELISA. (F) qRT–PCR detection of CCL5 expression levels in the sputum of patients infected with RSV. (G) CCL5 overexpression was verified in A549 cells transfected with pcAGGS-CCL5 by qRT–PCR. Following RSV infection, the intracellular viral protein levels in A549 cells transfected with pcAGGS-CCL5 were detected by Western blotting (H) or fluorescence microscopy (I). (J) After treatment with CCL5 protein, intracellular viral protein (GFP-tag) levels in A549 cells were detected by Western blotting. (Unpaired t test, \**P*<0.05, **P<0.01, \*\*\**P*<0.005, and \*\*\*\**P*<0.001)

Most studies on chemokines have focused on their recruitment of immune cells (such as monocytes/macrophages, granulocytes, natural killer [NK] cells and/or T cells) to enhance the immune response during inflammation and viral infection.

Following RSV infection, respiratory epithelial cells are highly induced to express CCL5, but the number of CCL5 receptors on respiratory epithelial cells is typically not increased. Does the interaction between CCL5 and its receptors on host cells impact RSV survival? To answer this question, after transfection with a eukaryotic plasmid expressing CCL5, A549 cells were infected with RSV and incubated for 24 hours. The results showed that with the increase in CCL5 expression level, the expression level of intracellular viral protein was significantly decreased (Fig. 1G-I). Therefore, it was suggested that the elevation in CCL5 expression may affect viral replication in respiratory epithelial cells. To test this hypothesis, we pretreated A549 cells with CCL5 protein for 30 min and then infected the cells with RSV (RSV-GFP-tag), and we found that GFP expression gradually decreased with increasing CCL5 concentration in the culture medium (Fig. 1J), indicating that the expression of CCL5 was negatively correlated with RSV replication.

### CCR1 overexpression affected the intracellular RSV load

CCL5 exerts its biological effect through its receptors, which include CCR1, CCR3 and CCR5. Therefore, we measured the expression levels of these three receptors in cells infected with RSV. We found that the expression level of CCR1 was decreased, that of CCR5 was increased, and that of CCR3 expression did not significantly change (Fig. 2A). Then, A549 cells were first transfected with CCR1, CCR3 or CCR5 recombinant expression plasmids (Fig. 2B) and then infected with RSV and incubated for 24 hours. Interestingly, the viral load (indicated by GFP [green] fluorescence) decreased significantly only after CCR1 was overexpressed (Fig. 2C and D). To determine whether CCR5 affects RSV load, CCR5 expression was knocked down by CCR5-specific short interfering RNA (siRNA), the effectiveness of which was verified by Western blotting (Fig. 2E). Subsequently, CCR5-knockdown cells were infected with RSV (MOI=0.5). The GFP viral tag was detected in cells through fluorescence microscopy and Western blot analysis (Fig. 2F and G), and the results showed no significant difference in RSV proliferation in the CCR5-knockdown cells and the normal controls. These data suggest that RSV proliferation in host cells was not affected by a change in the expression levels of CCR3 or CCR5 but was affected by changed CCR1 expression.

**Fig. 2.**
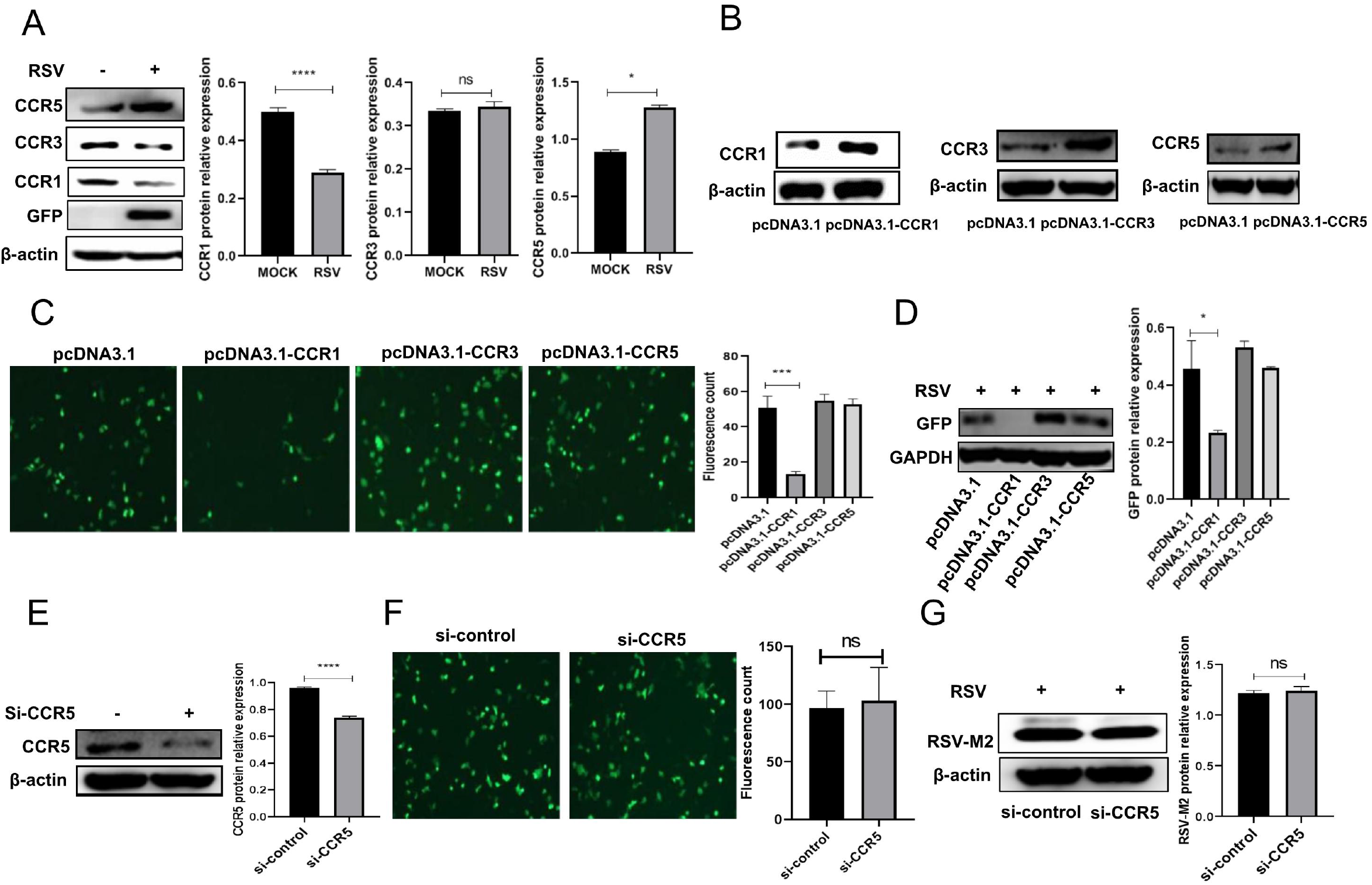
CCR1 expression decreased in A549 cells infected with RSV, and the CCR1 expression level was affected by the level of intracellular viral protein. (A) The expression of CCL5 receptors in A549 cells infected with RSV was analyzed by Western blotting. (B) A549 cells were transfected with 500 ng of CCR1, CCR3, CCR5 or an empty plasmid. Whole protein was extracted, and the transfection efficiency was verified by Western blotting. The viral load of A549 cells overexpressing CCR1, CCR3 or CCR5 was analyzed by fluorescence microscopy (C) and Western blotting (D). Western blot showing the effect of CCR5 gene knockout in A549 cells (E) that had been infected with RSV and incubated for 24 hours. The number of intracellular viruses was determined by fluorescence microscopy (F), and the level of intracellular viral proteins was detected by Western blotting (G). (Unpaired t test, \**P*<0.05, ***P<0.005, and \*\*\*\**P*<0.001)

### The CCR1 expression level was negatively correlated with RSV replication

After 24 hours of infection with RSV, viral proliferation was significantly reduced in A549 cells overexpressing CCR1 (1000 ng of pcDNA3.1-CCR1) compared with that in the control group cells (Fig. 3A and B), and with the increase in CCR1 overexpression, the intracellular viral load gradually decreased (Fig. 3C and D), suggesting that CCR1 expression was negatively correlated with RSV replication. Furthermore, A549 cells were pretreated with different concentrations of anti-CCR1 neutralizing antibodies and then infected with RSV, and the analysis showed that the viral load gradually recovered in a dose-dependent manner with the increase in neutralizing antibody concentration (Fig. 3E and F), indicating that RSV replication was resumed after the binding of CCR1 and its ligand was blocked. Then, after RSV infection of CCR1-knockdown A549 cells for 24 hours, the intracellular viral protein level was increased significantly compared with that in the control cells (Fig. 3G), confirming that the expression level of CCR1 was negatively correlated with the intracellular replication of RSV.

**Fig. 3.**
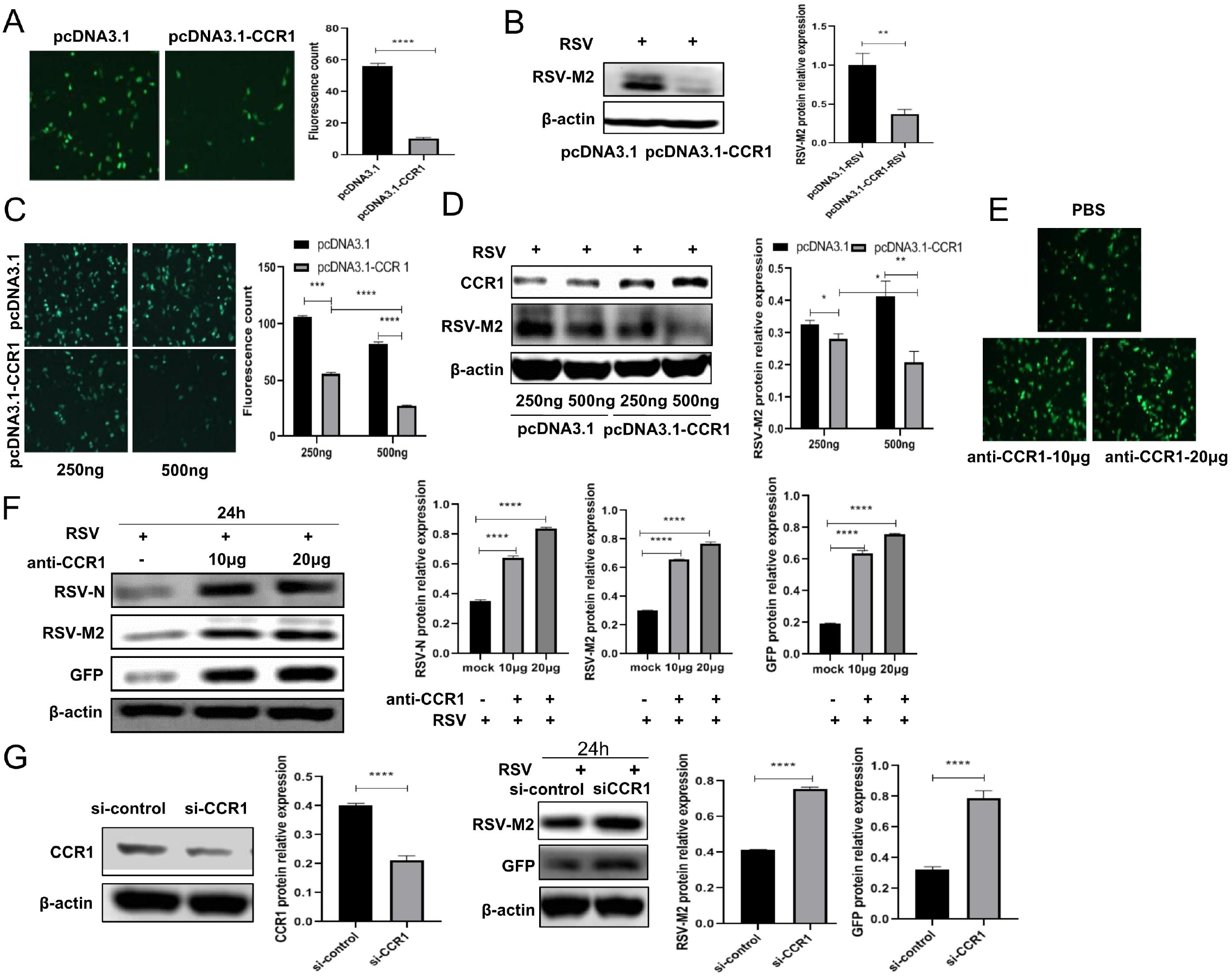
CCR1 expression was negatively correlated with RSV replication. (A. B) The viral load in A549 cells overexpressing CCR1 was analyzed by fluorescence microscopy and Western blotting. (C, D) The relationship between CCR1 overexpression and intracellular viral load was analyzed by fluorescence microscopy and Western blotting. (E, F) The effect of the anti-CCR1 neutralizing antibody on RSV intracellular replication was measured by fluorescence microscopy and Western blotting. (G) Western blot showing the RSV protein expression in CCR1-knockdown A549 cells following RSV infection for 24 hours. (Unpaired t test, \**P*<0.05, **P<0.01, \*\*\**P*<0.005, and \*\*\*\**P*<0.001)

### The obvious decrease in viral load after CCR1 was overexpressed was not caused by up-regulating type I interferon level, inducing cell apoptosis, down-regulating the expression of CX3CR1, or inhibiting RSV adhesion and entry into cells

How does CCR1, a membrane receptor, affect virus replication? To answer this question, after the CCR1 expression plasmid (500 ng) was transfected into A549 cells, we measured CCR1 expression through confocal microscopy (Fig. 4A) and Western blotting of extracted membrane proteins (Fig. 4B). We found showed that CCR1 expression on the membrane surface had significantly increased, indicating that CCR1 overexpression in A549 cells enhanced the interaction between CCR1 and its ligand CCR1. What is the mechanism of RSV load reduction following CCR1 overexpression? Does CCR1 overexpression increase IFN-I expression to enhance cellular antiviral function, or does it lead to apoptosis and the subsequent decrease in intracellular viral load? CCR1 overexpression affect virus adhesion, entry into cells, intracellular proliferation and/or budding? In answering these questions, we found that compared with that in the control group, the IFN-I expression level induced by CCR1-overexpressing cells was not increased or decreased, suggesting that the decrease in intracellular viral load in CCR1-overexpressing cells was not a result of increased IFN-I pathway activity (Fig. 4C). Additionally, A549 cells overexpressing CCR1 showed no obvious change in apoptosis rate regardless of their RSV infection status, suggesting that the decrease in viral load following CCR1 overexpression was not caused by apoptosis (Fig. 4D). RSV was incubated with cells at 4°C until it adhered to cells, and then, the RSV-attached cells were cultured at 37°C for 30 min to initiate virus entry into these cells. Then, the cells were collected, and the degree of viral adhesion and entry into cells was determined by real-time PCR. The results showed that the viral load in cells overexpressing CCR1 was not significantly different than that of the control group cells, suggesting that CCR1 overexpression did not affect viral adhesion or entry into cells (Fig. 4E). Simultaneously, the expression of CX3CR1, the main receptor of RSV, was detected in CCR1-overexpressing A549 cells by Western blotting and confocal microscopy, and the results showed that the CX3CR1 expression level did not significantly change compared with that in the control cells (Fig. 4F and G). Thus, the decrease in intracellular viral load caused by CCR1 overexpression was not due to an effect on the level of the main RSV receptor. These results showed that CCR1 overexpression had almost no influence on cell apoptosis; the expression of IFN-I or CX3CR1; or RSV adhesion or entry into cells.

**Fig. 4.**
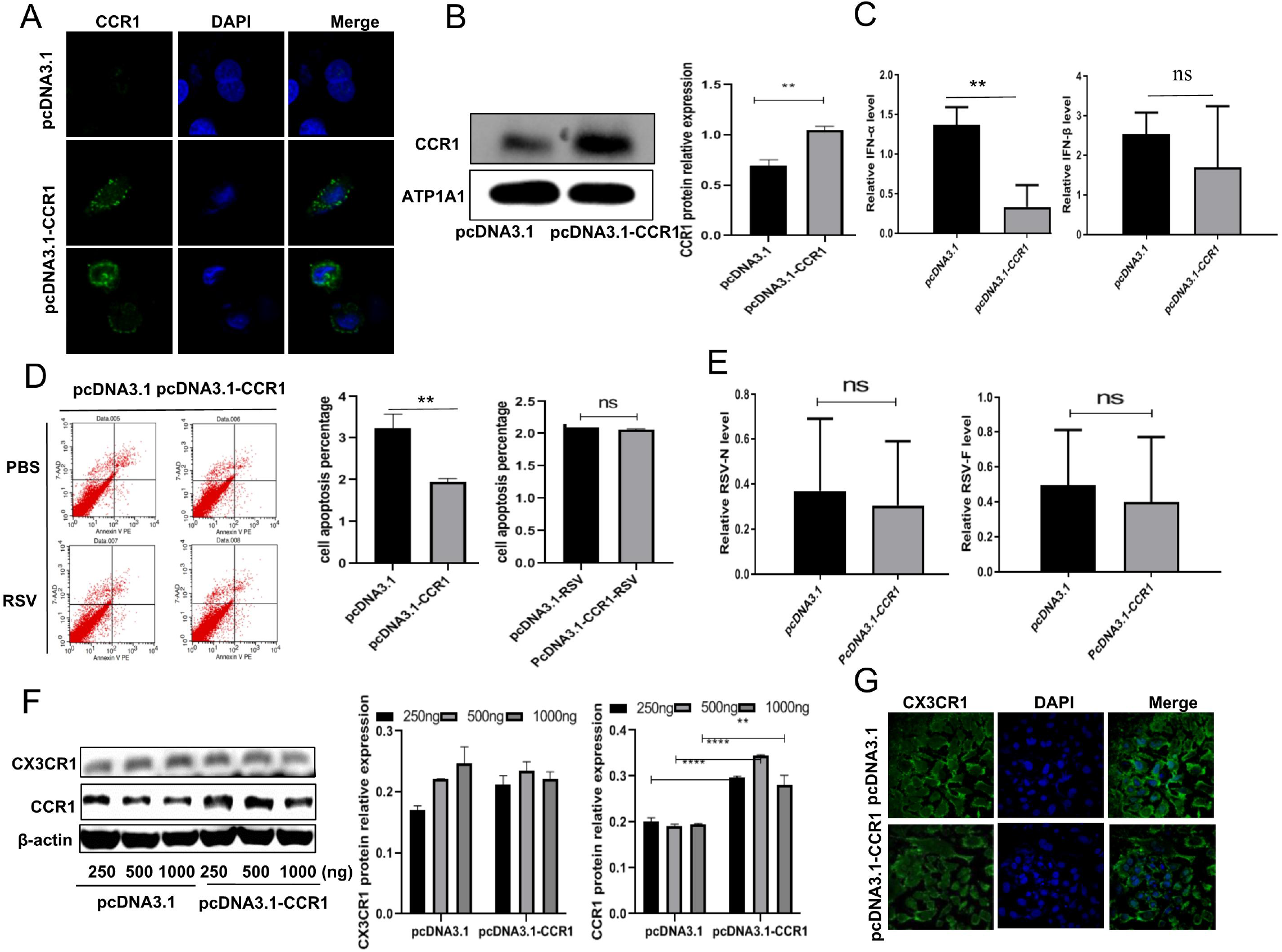
Possible factors leading to the decrease in viral load following CCR1 overexpression. The localization of overexpressed CCR1 in A549 cells was analyzed by confocal microscopy (A) and Western blotting (B). After RSV infection, the effect of CCR1 overexpression on the level of type I interferon was detected by qRT–PCR (C), and cell apoptosis was induced and measured by flow cytometry (D). (E) The effect of CCR1 overexpression on RSV adhesion and entry was detected by qRT–PCR. Western blotting (F) and confocal microscopy (G) were performed to investigate the effect of CCR1 overexpression on CX3CR1 expression. (Unpaired t test, \*\**P*<0.01 and \*\*\*\**P*<0.001)

### CCR1 suppresses RSV replication by downregulating the activity of HSP90 (an RSV polymerase chaperone)

It has been reported that the RSV RNA polymerase complex consists of viral nucleoprotein (N), large polymerase protein (L) and its cofactor, phosphate protein (P). The M2-1 protein (which is a polymerase cofactor) is also required for transcriptional activity. In addition, host cell proteins HSP90 and HSP70 have been shown to play an essential role in the RNA synthesis of many viruses, and in particular, HSP70 has been found to play a direct role in RSV polymerase activity in human RSV-infected cells. HSP90 plays an important role in the stability of RSV polymerase. CDC2, a known client protein of HSP90, is often used as a marker of HSP90 activity. In this study, we found that after the RSV infection of A549 cells, the CDC2 protein level increased, and HSP70 expression did not change significantly, suggesting that following RSV infection, the activity of the CDC2 partner, HSP90, may have been increased, promoting RSV replication (Fig. 5A). Furthermore, after blocking CCR1 overexpression in A549 cells with different concentrations of neutralizing antibodies against CCR1, the cells were infected with RSV and incubated for 24 hours. The results showed that the RSV protein expression level increased gradually with the increase in CCR1 neutralizing antibody, suggesting that blocking the binding of CCR1 to its ligand restores RSV replication. Moreover, the expression levels of HSP90 and CDC2 decreased after the CCR1-overexpressing cells were infected with RSV. In contrast, with the increase in the anti-CCR1 neutralizing antibody, the levels of CDC2 and viral proteins increased gradually, although the protein level of HSP70 did not change significantly (Fig. 5B), suggesting that CCR1 exerted a negative regulatory effect on HSP90 activity. In addition, after the binding of CCL5 and CCR1 was blocked, RSV infection induced HSP90 activity in A549 cells to promote viral replication, which may explain the downregulation in CCR1 protein expression downregulated after RSV infection (Fig. 2A). Tanespimycin (17-AAG) is an effective inhibitor of HSP90 activity. After optimizing the concentration of 17-AAG with a cell counting kit-8 (CCK-8) assay kit, A549 cells were treated with 17-AAG, and then, with increases in 17-AAG concentration, the CDC2 protein level was found to be gradually decreased in a dose-dependent manner (Fig. 5C). Then, following pretreatment with 17-AAG, the cells were infected with RSV. With the increase in inhibitor concentration, the level of intracellular viral protein also gradually decreased, indicating that HSP90 activity affects intracellular RSV replication (Fig. 5D and E). To confirm that CCR1 activation by its ligand CCL5 affects RSV intracellular replication, CCR1-overexpressing cells were treated with CCL5, and the results showed that the CDC2 protein level decreased gradually with increasing CCL5 protein concentration (Fig. 5F), suggesting that CCR1 activation by its ligand CCL5 downregulates HSP90 activity. In contrast, the level of intracellular viral protein or GFP in the knockdown cells was significantly lower than that in the control cells (Fig. 5G-I). Therefore, we speculated that CCR1 binding to its ligand CCL5 can downregulate HSP90 activity, which in turn affects RSV replication.

**Fig. 5.**
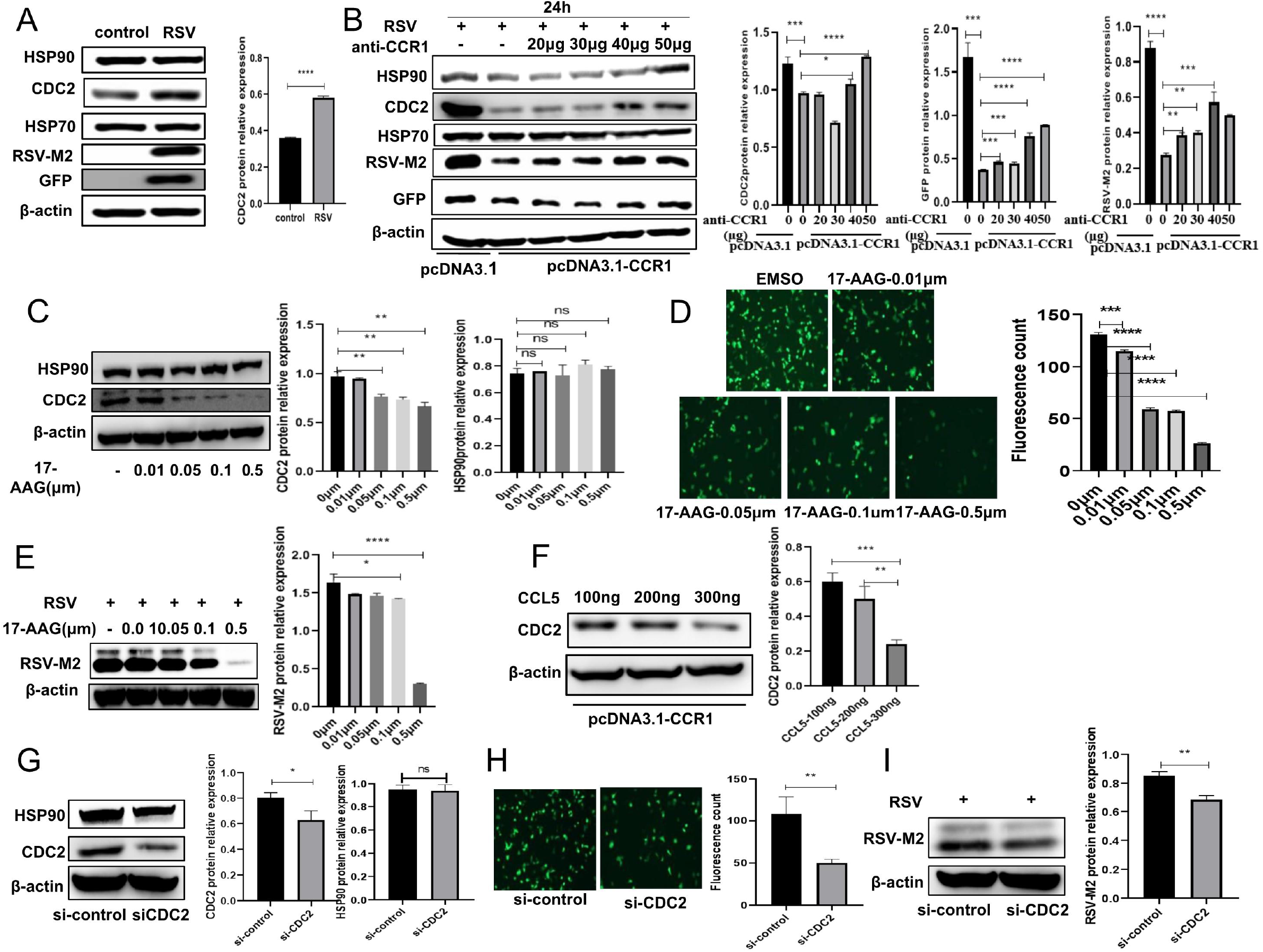
Analysis of HSP90 activity affecting the intracellular replication of RSV. (A) The effect of RSV infection on the expression of an RSV chaperone protein was analyzed by Western blotting. (B) The effects of CCR1 overexpression and the anti-CCR1 neutralization antibody on CDC protein levels (representing the activity of HSP90) in A549 cells infected with RSV were detected by Western blot. (C) Western blot analysis of CDC2 protein levels in A549 cells treated with the HSP90 inhibitor 17-AAG. The effect of 17-AAG on RSV intracellular replication was analyzed by fluorescence microscopy (D) and Western blotting (E). (F) After CCR1-overexpressing A549 cells were treated with different concentrations of the CCL5 protein, the CDC2 protein level was detected by Western blotting. (G) The effect of CDC2 knockdown in A549 cells was verified by Western blotting. The effect of CDC2 knockdown on RSV replication in A549 cells was analyzed by fluorescence microscopy (H) and Western blotting (I). (Unpaired t test, \**P*<0.05, \*\**P*<0.01, \*\*\**P*<0.005, and \*\*\*\**P*<0.001)

### Activation of CCR1 by CCL5 leads to the preferential recruitment of the adaptor Gαi, which regulates HSP90 activity to affect RSV replication

To further study the possible molecular mechanism by which CCR1 inhibits HSP90 activity, we needed understand the process through which the chemokine receptor CCR1 is activated. CCR1 belongs to the seven transmembrane G protein-coupled receptor (GPCR) family. The activation of GPCRs depends on agonists in the extracellular environment, particularly those that preferentially recognize intracellular effectors to induce signalling farther downstream. The most common effectors in cells are adaptor G proteins or β-arrestin (β-arr)(10). Therefore, in this study, A549 cells were cotransfected with CCR1, Gαi or β-arrestin2 recombinant plasmids for 24 hours and then stimulated with CCL5 protein. Through confocal microscopy, we observed that CCR1 colocalized with the adaptors Gαi and β-arrestin2 following CCL5 treatment (Supplementary Fig. A and B). However, under the same conditions, we performed co-immunoprecipitation (co-IP) and found that CCR1 derived from transfected A549 cells bound more Gαi and HSP90 and less β-arrestin 2 than the CCR1 derived from CCL5-deficient control cells (Fig. 6A and B), suggesting that CCR1 activated by CCL5 preferentially recruits Gαi not β-arrestin 2. As expected, when Gαi was overexpressed or knocked down, Gαi was negatively correlated with CDC2 protein level (which represents HSP90 activity) after RSV stimulation (Fig. 6C and D), and a similar result was obtained after CCL5 protein treatment (Fig. 6E). Interestingly, the ERK phosphorylation level was not related to Gαi but was positively correlated with β-arrestin 2 (Fig. 6C). All these results indicated that activated CCR1 preferentially recruits Gαi to inhibit RSV replication by regulating HSP90 activation in a mechanism independent of the ERK pathway. Although another adaptor, β-arrestin2, might promote HSP90 activity by activating ERK kinase, β-arrestin 2 was rarely recruited by CCL5-activated CCR1.

**Fig. 6.**
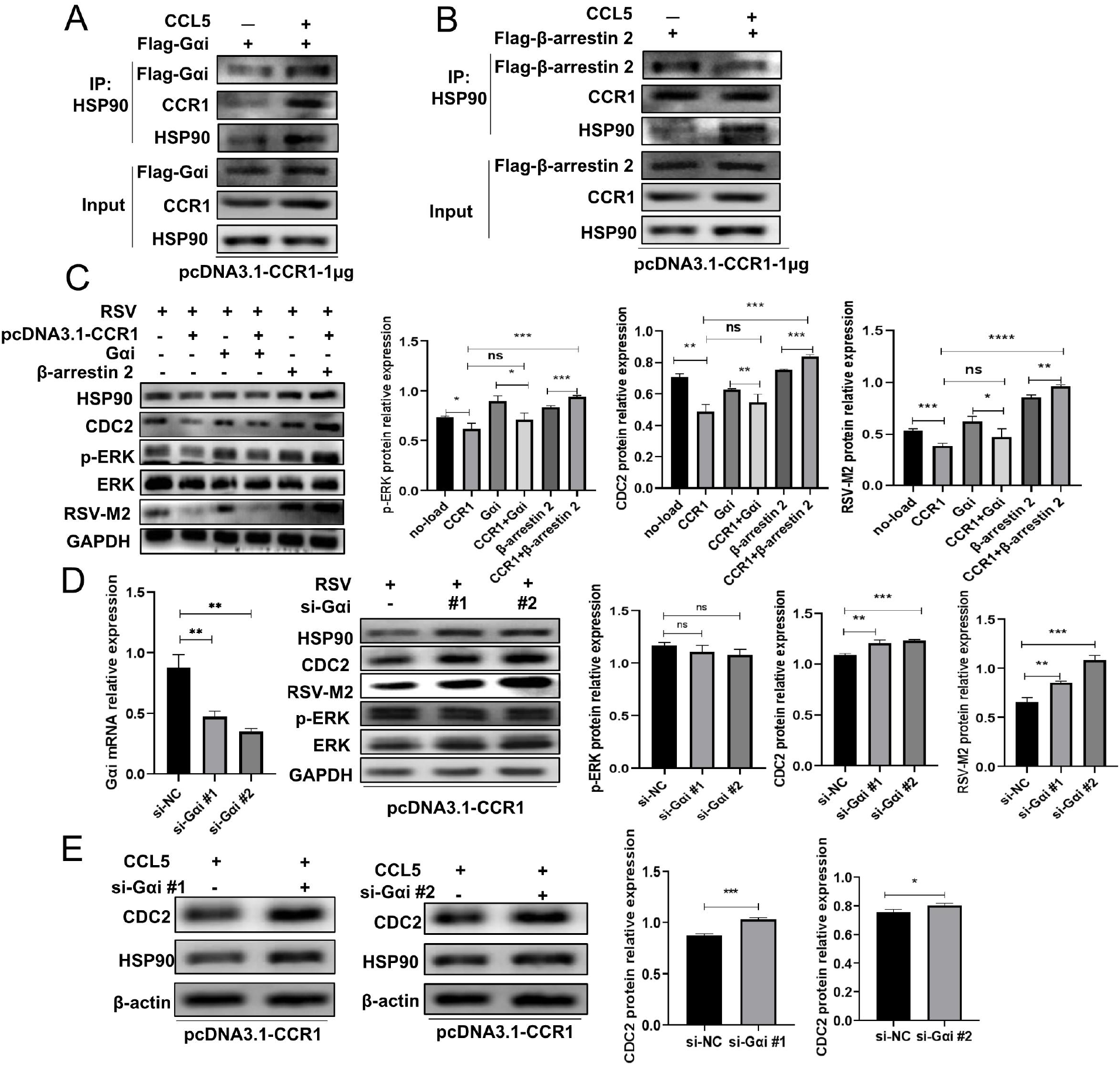
Gαi negatively regulates HSP90 activity after recruitment and activation. After overexpression of CCR1 and Gαi or β-arrestin2, the interaction between HSP90 and Gαi or β-arrestin2 in A549 cells treated with CCL5 was detected by co-IP (A, B), or the expression levels of CDC2, p-ERK and RSV-M2 in A549 cells stimulated with RSV were detected by Western blotting (C). Following CCR1 overexpression and Gαi knockdown, A549 cells were infected with RSV, and the expression levels of CDC2, p-ERK and RSV-M2 were detected (D), or A549 cells were treated with CCL5, and the expression level of CDC2 was detected (E) by Western blotting. (Unpaired t test, \**P*<0.05, \*\**P*<0.01, \*\*\**P*<0.005, and \*\*\*\**P*<0.001)

### Activated Gαi upregulates inositol 1,4,5-triphosphate receptor type 3 (ITPR3) expression, which inhibits STAT3 phosphorylation, thereby affecting HSP90 activity and viral replication

To identify the signaling pathway by which Gαi affects HSP90 activity, we analyzed differentially expressed proteins with liquid chromatography with tandem mass spectroscopy (LC–MS/MS). On the basis of their expression in the control group, 58 differentially expressed proteins were identified in the CCL5-treatment group (Fig. 7A); among these proteins, the information on 12 corresponding genes was unclear (not analyzed). The remaining 46 genes were subjected to GO enrichment analysis, and the results of GO molecular function analysis showed that most of these genes were associated with RNA binding (Fig. 7B). In addition, a protein–protein interaction (PPI) analysis was performed with the remaining 46 genes, and then, a protein interaction network was constructed by removing irrelevant genes and adding certain genes that might show potential protein interactions (Fig. 7C), and on this basis, we identified the CCR1, ITPR3, SFPQ and GRB2 genes as encoding proteins with potential Gαi interactions. GRB2 was not found in the LC–MS/MS results; SFPQ was not enriched in any pathway in the Kyoto Encyclopedia of Genes and Genomes (KEGG) enrichment analysis; and neither ITPR3, CCL5, CCR1 nor Gαi was found to participate in the HSA05163 (human cytomegalovirus infection) signaling pathway. Therefore, we focused on the ITPR3 protein. In A549 cells overexpressing CCR1, we knocked down ITPR3 expression and then treated the cells with CCL5 protein or infected them with RSV. The results showed that the expression level of CDC2 was significantly enhanced compared with that in the control group, and the viral protein level was significantly increased in the RSV-infected group (Fig. 7D and E), suggesting that activation of Gαi can downregulate HSP90 activity by upregulating ITPR3 and can inhibit RSV replication. These results encouraged us to further analyze the ITPR3 protein interactions, and upon examining LC–MS/MS results, we found a close correlation between ITPR3 and STAT3 expression (Fig. 7C). Therefore, we re-examined the CCR1-overexpressing cells infected with RSV. We were excited to find that the phosphorylation level of STAT3 was significantly reduced in these cells compared with that in the control cells (Fig. 7F), and following knockdown of ITPR3 expression, both the phosphorylation level of STAT3 and the activity of HSP90 were significantly increased under the same conditions (Fig. 7G and H), suggesting that STAT3 may be a downstream molecule involved in the negative regulation of ITPR3. When an inhibitor of STAT3 (stattic) inhibited STAT3 phosphorylation, the activity of HSP90 was significantly inhibited, but the expression level of ITPR3 was not affected, indicating that STAT3 is a downstream molecule that is negatively regulated by ITPR3 and that HSP90 is the downstream molecule of STAT3 (Fig. 7I). Thus, we demonstrated that activated CCR1 tends to preferentially recruit Gαi and that Gαi upregulates ITPR3 expression to inhibit STAT3 phosphorylation (Fig. 7I and J), regulating the activity of RSV RNA polymerase chaperone HSP90 to modulate viral replication.

**Fig. 7.**
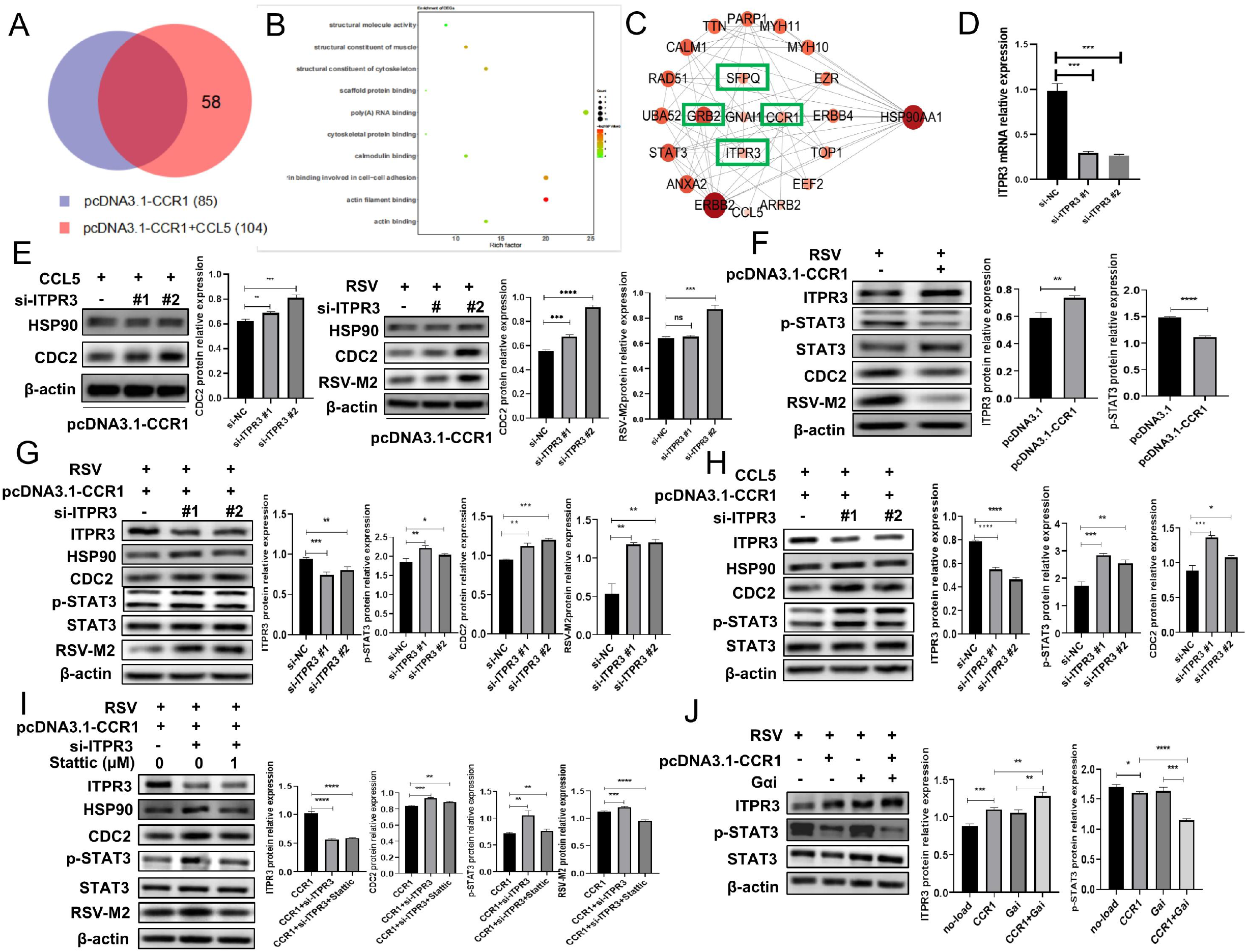
Gαi recruited by activated CCR1 upregulates ITPR3, which inhibits STAT3 phosphorylation, thereby affecting HSP90 activity and viral replication. After CCR1-overexpressing A549 cells were treated with or without CCL5, mass spectrometry was performed to analyze the differentially expressed proteins between the two groups (A), and GO enrichment with differentially expressed genes (B) and the interactions of proteins encoded by differentially expressed genes (C) were analyzed. (D) The effect of knocking down ITPR3 expression was verified by qRT–PCR. (E) After ITPR3 expression was knocked down, CCR1-overexpressing A549 cells were stimulated with CCL5 or RSV, and the protein expression levels of CDC2 and RSV-M2 were detected by Western blotting. (F) Following RSV infection, the expression of ITPR3, p-STAT3, CDC2 and viral proteins in CCR1-overexpressing cells was determined by Western blotting. (G, H) After ITPR3 expression was knocked down, CCR1-overexpressing A549 cells were stimulated with RSV or CCL5, and the protein expression level of p-STAT3 was detected by Western blotting. (I) After ITPR3 expression was knocked down, A549 cells overexpressing CCR1 were treated with a STAT3 inhibitor (stattic) and then stimulated with RSV for 24 hours. The protein expression levels of CDC2, p-STAT3 and RSV-M2 were detected by Western blotting. (J) After A549 cells overexpressing CCR1 and/or Gαi were infected with RSV, the protein expression of ITPR3 and p-STAT3 was detected by Western blotting. (Unpaired t test, \**P*<0.05, \*\**P*<0.01, \*\*\**P*<0.005, and \*\*\*\**P*<0.001)

### CCR1 overexpression in respiratory epithelial cells was confirmed to inhibit RSV replication in vivo

To confirm the role of CCR1 in regulating RSV replication in vivo, BALB/c mice were challenged with RSV, and then, the expression of CCL5 and CCR1 was evaluated. Seventy-two hours after RSV infection, the CCR1 level was significantly decreased compared with that in the mock group (Fig. 8B), although the CCL5 expression level in bronchoalveolar fluid (BALF) was much higher than that in the mock group (Fig. 8A). These data were consistent with those obtained at the cellular level, indicating that RSV may inhibit the antiviral ability of host cells by downregulating CCR1 level to facilitate its own survival in vivo. Therefore, we constructed an adeno-associated virus (AAV) carrying the CCR1 sequence (AAV-CCR1), which are specifically expressed in lung epithelial cells. BALB/c mice were infected with the recombinant virus (intranasally [i.n.]) to overexpress CCR1 in lung epithelial cells. After some mice were sacrificed to determine the expression of the AAV marker RFP and CCR1 in the lung (Fig. 8C and D), the remaining mice were challenged with RSV. The expression of CCR1 and replication of RSV were measured, and lung inflammation was analyzed. Notably, the lung RSV viral load was markedly reduced in the CCR1-overexpressing mice, as detected by qRT–PCR (Fig. 8E) or whole-lung imaging, which showed weaker fluorescence (GFP) in the CCR1-overexpressing mice infected with RSV than in the control mice (Fig. 8F), indicating that CCR1 overexpression inhibited RSV replication in vivo. In addition, lung inflammation in the CCR1-overexpressing mice was significantly reduced compared with that in the control group mice. In detail, the pathological manifestations indicated a mild collapse of the alveolar cavity, mild proliferation of alveolar cells, and a reduction in focal lymphocyte aggregation and monocyte/phagocyte infiltration in the alveolar cavity (Fig. 8G), suggesting that in the early stage of infection, the greater the pulmonary RSV load is, the more obvious the pulmonary inflammation.

**Fig. 8.**
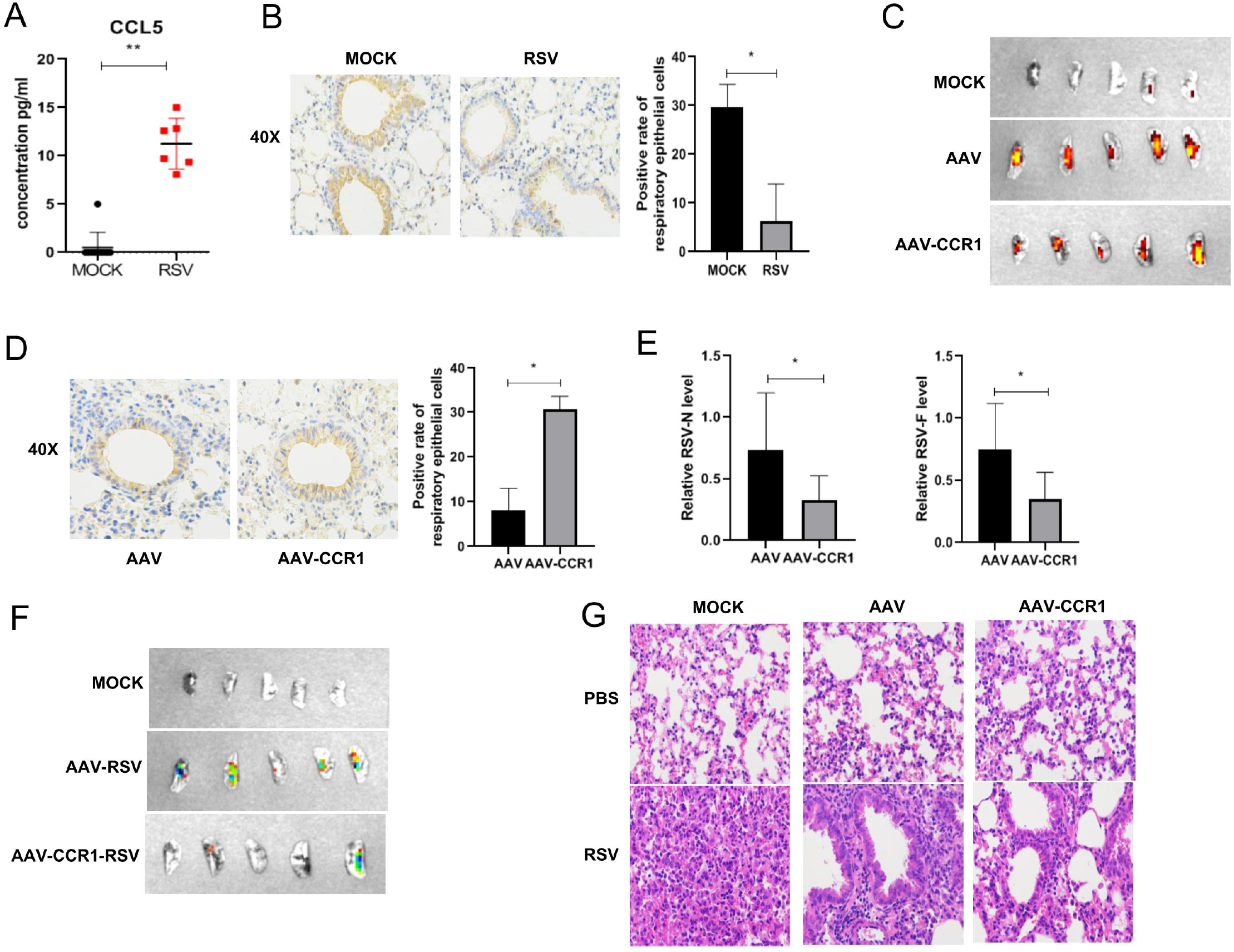
Analysis of the effect of the chemokine receptor CCR1 binding to its ligand in an animal model. (A) CCL5 expression in the BALF of RSV-infected mice was detected by qRT–PCR. (B) CCR1 expression in the respiratory epithelial cells of RSV-infected mice was analyzed by immunohistochemistry. (C) RFP expression in the lungs of AAV- or AAV-CCR1-transfected mice was detected by whole-lung imaging. (D) CCR1 expression in the lungs of AAV- or AAV-CCR1-transfected mice was detected by immunohistochemistry. (E) The mRNA expression levels of the RSV N and M genes in lung tissue of AAV- or AAV-CCR1-transfected mice were detected by qRT–PCR. (F) The RSV load was detected in the lungs of AAV- or AAV-CCR1-transfected mice by whole-lung imaging. (G) Lung inflammation was analyzed by H&E staining. (Unpaired t test, \**P*<0.05 and \*\**P*<0.01)

## DISCUSSION

Viral infection is one of the major causes of human morbidity and mortality worldwide. In particular, respiratory viruses are the main pathogens causing global pandemics, posing a serious threat to human health and imposing a heavy economic burden on society. Increasing evidence has shown that the occurrence and development of respiratory tract infections are closely related to the host immune defense system. The innate immune system is based on both constitutive and inducible mechanisms and eliminates infections and attenuates self-damage to maintain homeostasis. Inducible mechanisms such as those mediated through pattern recognition receptors (PRRs), as well as antigen-specific receptors, are activated only in response to stimuli and show the ability to induce very strong and efficient immune responses but can also lead to inflammation(11). The types of inflammation and the development and prognosis of infection-induced diseases are profoundly affected by the interaction between viruses and immune molecules, of which chemokines and their receptors are important components. Inflammatory cells are the main sources and targets of chemokines. However, it has been reported that some nonimmune cells, such as respiratory epithelial cells, can produce chemokines or respond to certain chemokines, especially after exposure to pathogens(12,13). In this study, we confirmed that RSV induced high expression of the chemokine CCL5 in respiratory epithelial cells in vivo and in vitro. Specifically, sputum samples of children with clinical respiratory tract infection (0.5-13 years old) were collected, and multiple detection kits were used to detect 13 kinds of respiratory pathogens. Finally, 32 patients with simple RSV infection and 66 negative control samples were selected for detecting the relative expression level of CCL5 mRNA by qRT–PCR. The analysis showed that the CCL5 expression level in most of the clinical RSV-infected samples was much higher than that in the negative control samples. However, the expression level of some RSV-infection samples was similar to that of the negative control samples; this outcome was due to the self-limited expression of cytokines in convalescent patients or individual differences (Fig. 1F).

The interaction between human host cells and viruses is realized through a complex mechanism in which cytokines play important roles. Among the large group of cytokines, chemokines are currently considered to be among the essential to immunopathology and/or defense against viral infection. Chemokines typically bind to their receptors to trigger a variety of biological effects: 1. Chemokines determine the type of inflammation that is manifested by recruiting and interacting with inflammatory cells. For example, CCR5 and CXCR3 are mainly expressed in activated T cells and are related to the Th1 immune response, while CCR3 and CCR4 are related to the Th2 response. 2. Some chemokine receptors are receptors or coreceptors of viruses, and therefore, their expression is the determinate for whether viruses can invade host cells(14-16). In this study, we found that chemokines bound to receptors on the surface of host cells to regulate the response of host cells to viruses and then affected the replication of intracellular viruses. Receptors that can bind to CCL5 include CCR1, CCR3 and CCR5, which are expressed not only on the surface of immune cells but also on some nonimmune host cells, such as respiratory epithelial cells. When we infected A549 cells with RSV, we found that CCR1 expression was downregulated, and a similar decrease in CCR1 protein levels has been reported following coronavirus infection(17). The expression of CCR3 or CCR5 was upregulated or unchanged, but the viral load increased with prolonged infection time (Fig. 2A), suggesting that the three aforementioned receptors may have distinct functions after being activated by CCL5 in RSV infection. Interestingly, our results showed that the RSV load decreased significantly only in cells overexpressing CCR1 when all three receptors were overexpressed in A549 cells (Fig. 2B-D). That is neither CCR3 nor CCR5 exerted an effect on viral load, indicating that activated CCR1 may be involved in regulating the intracellular RSV load. What is the mechanism by which CCR1 regulates viral load? To answer this question, we first confirmed that CCR1 overexpression on the cell surface did not affect the protein level of CXCR3 (a receptor known for enabling RSV entry into a cell). Second, we measured viral adhesion and entry into CCR1-overexpressing cells. The results indicated that overexpressing CCR1 did not affect viral adhesion and entry into cells. Third, we measured the apoptosis rate and IFN-I expression in A549 cells overexpressing CCR1 after RSV infection, and we found that overexpression of CCR1 on these cells neither increased IFN-I production nor the apoptosis rate (Fig. 4). These results did not clarify the mechanism by which CCR1 overexpression on the cell membrane led to the significant reduction in intracellular viral load.

Since the genome of a virus is very small, viruses mainly rely on the biosynthetic components of the host cell to replicate. We analyzed previously obtained transcriptome data on A549 cells infected with RSV(18) and found that three host proteins, HSP90, HSP70, and STIP1, were highly expressed, which was consistent with the results reported by Munday et al.(19). After securing experimental verification, we found that, among these proteins, HSP90 was the most meaningful to our study. HSP90 is an essential molecular chaperone required for the maintenance, activation and/or maturation of specific client proteins. HSP90 is an ATPase and is expressed as two subtypes, HSP90α and HSP90β, usually in the cytoplasm as homodimers. The structure of each monomer includes the C-terminal dimerization domain, intermediate domain and structurally unique N-terminal ATPase domain(20). Through changes in its conformation, HSP90 regulates the activity, maturation, localization and turnover of many selective substrate proteins, which are called HSP90 clients(21). Early biochemical studies have shown that the binding of ATP regulates the conformational transition of HSP90, which plays a crucial role in recognizing and activating client proteins. Before ATP binding, HSP90 is in a V-shaped conformation (open state), and after ATP binding, a series of conformational changes are triggered, resulting in a more condensed N-terminal dimerization domain (closed state). Cyclin-dependent kinase 1 (CDK1) (also known as the cell division control protein CDC2) is an HSP90 client protein that depends on the stable chaperone activity of HSP90, and therefore, CDK1 (CDC2) is often used as a marker of HSP90 activity, especially when HSP90 inhibitors are unavailable^19^. In fact, the replication of many viruses, including DNA viruses, positive and negative RNA viruses and double-stranded RNA viruses, almost universally depends on HSP90,. Most of the identified viral client proteins of HSP90 are capsid proteins and viral polymerases(22). A variety of respiratory virus polymerases are HSP90-dependent, including RNA polymerases of positive and negative RNA viruses and DNA-dependent DNA polymerases, such as those of vesicular stomatitis virus (VSV) and other paramyxoviruses(23), Ebola virus(24) and influenza virus(25), although these proteins differ in terms of evolution, structure and function(26). The polymerase activity of distinct viruses relies on the auxiliary effect of HSP90 chaperone molecules, indicating that HSP90 may recognize the conserved characteristics of these different enzymes or may promote a common step in their maturation and/or replication, such as interactions with nucleic acids. Our study also showed that the CDK1 (CDC2) protein level, which represents the activity of HSP90, was increased after RSV infection. When the activity of HSP90 was suppressed by 17-AAG (an inhibitor of HSP90), the RSV load and RSV replication in A549 cells were also significantly suppressed (Fig. 5C-E). Munday et al.(19) reported that HSP90 mainly interacts with the L protein of RSV polymerase to promote viral replication. These results imply that HSP90 plays an essential role in the lifecycle of RSV.

When CCR1 was overexpressed in A549 cells that were subsequently infected with RSV, we found that the CDC2 protein level was significantly reduced. Then, we blocked CCR1 activity with an anti-CCR1 neutralization antibody and infected the cells with RSV. We were subsequently surprised to find that with increasing concentrations of neutralizing antibody, the protein levels of HSP90 and CDC2 gradually recovered, and the expression of RSV viral protein increased in a dose-dependent manner (Fig. 5B), indicating that CCR1 might negatively regulate HSP90 activity and viral replication.

What is the molecular mechanism by which CCR1 affects HSP90 activity? This chemokine receptor belongs to the GTP protein-coupled transmembrane receptor family, and it is composed of approximately 330 amino acids. It contains seven transmembrane regions that divide the molecule into several domains: the N-terminal domain, 3 extracellular rings, 3 intracellular rings and the C-terminal domain(27). The effect of activated GPCR depends on the agonists in the extracellular environment and the intracellular effectors that are recognized preferentially to induce signals farther downstream signals. The most common intracellular effector is a G protein or β– arrestin(10). In a common feature of chemokine binding to receptors, the intracellular helices of the activated receptor move outward, especially the transmembrane (TM)-VI and -VII helices, to make room for subsequent G protein and/or β–arrestin binding(28,29). In this study, we found that Gαi was mainly recruited by CCL5-activated CCR1, and further analysis revealed that, by recruiting different effectors, the chemokine receptor CCR1 induced different intracellular signaling pathways to produce different effects. It has been reported that β-arrestin 2 is recruited to induce ERK1/2 phosphorylation by activating the mitogen-activated protein kinase (MAPK) pathway, which then activates a series of downstream effector proteins and regulates the activity of diverse proteins(18). In our study, CCR1 and β-arrestin 2 were overexpressed in A549 cells that were then infected with RSV, and the levels of ERK phosphorylation and CDC2 and viral protein expression were found to be significantly higher in the cells overexpressing both CCR1 and β-arrestin2 than the cells overexpressing only CCR1 (Fig. 6C). When we inhibited ERK phosphorylation with inhibitors, the CDC2 level gradually decreased with increasing inhibitor concentration, and viral replication gradually decreased (Supplementary Fig. C). Therefore, we speculated that β-arrestin is recruited and subsequently activates ERK1/2, which upregulates HSP90 activity by influencing the expression or function of a cochaperone, thereby regulating the ability of the cochaperone to facilitate viral replication. However, we found that CCL5-activated CCR1 preferentially recruited Gαi, which rarely affects the ERK kinase pathway (Fig. 6C and D) and that it inhibited STAT3 phosphorylation by upregulating ITPR3, which is essential for both immune activation and cancer pathogenesis. Our further analysis revealed that STAT3 is a crucial kinase that promotes HSP90 activity. Considering these results, we verified that CCL5-activated CCR1 reduced HSP90 activity and inhibited the intracellular replication of RSV mainly through the Gαi/ITPR3/STAT3 signaling pathway.

In addition, we utilized tissue-specific promoter AAV technology to specifically overexpress CCR1 in the lung epithelial cells of mice, which were then infected with RSV. Within three days of infection, before the virus activated the specific immune response, the animals were sacrificed, and the viral load was found to be significantly decreased (Fig. 8E and F) compared with that of the control group mice.

Taken together, our results revealed a novel mechanism by which RSV escapes innate immunity. That is, although it induces high CCL5 expression, RSV might attenuate binding CCL5 by downregulating the expression of CCR1 in respiratory epithelial cells to weaken the inhibitory effect of CCR1 on HSP90 activity and thereby facilitate RSV replication in nonimmune cells (Fig. 9). Interestingly, we found similar results in CCR1-overexpressing cells infected with influenza virus (data not shown). To determine whether the mechanism we describe is common in pathogenic infection, further study is needed.

**Fig. 9.**
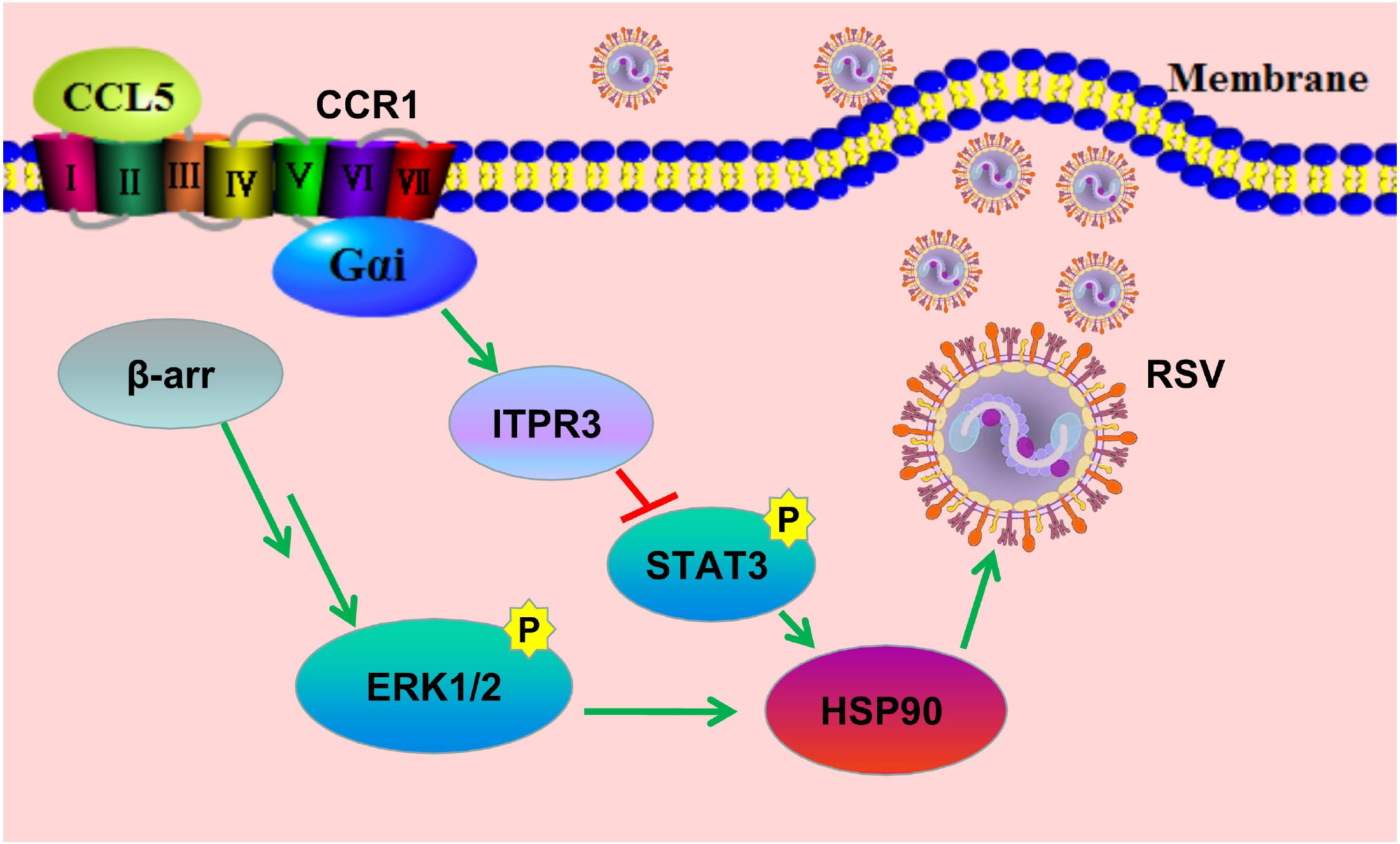
Schematic diagram showing activated CCR1 downregulating HSP90 activity through the Gαu/ITPR3/p-STAT3 pathway in respiratory epithelial cells. The chemokine CCL5 binds with CCR1 to downregulate HSP90 activity through the Gαi/ITPR3/p-STAT3 pathway, affecting the function of RSV RNA polymerase and inhibiting viral replication.

## ONLINE METHODS

### Patients

This study was conducted in accordance with the principles of the Declaration of Helsinki. Between 19 September 2018 and 25 February 2019, sputum specimens of pediatric inpatients were randomly collected with a sputum aspirator based on National Clinical Laboratory Procedures. The specimens were examined using a multiple pathogen detection kit designed to identify 13 respiratory pathogens (Health Gene Tech, Ningbo, China) in the Second Hospital of Hebei Medical University. Total RNA was collected and qualified according to the kit instructions, and 13 common respiratory pathogens were detected in the sputum samples by qRT–PCR and capillary electrophoresis. These pathogens were influenza A virus H1N1 and H3N2, parainfluenza virus species, human metapneumovirus, influenza B virus species, RSV, coronavirus, rhinovirus, bocavirus, *Chlamydia, Mycoplasma pneumoniae*, and adenovirus species. Sixty-six samples were selected as negative controls, and in these samples, none of these pathogens were found. For the experimental group, 32 samples were selected, and among the 13 pathogens identifiable with the kit, only RSV was found. Written informed consent was obtained from all parents or guardians.

### Animals and treatment

Four-week-old female specific-pathogen-free (SPF) BALB/c mice were purchased from the Experimental Animal Center of Hebei Medical University. All the mice were housed in temperature-controlled individual ventilated cages (IVCs) with a 12-hour light/12-hour dark cycle and were fed standard chow and sterile tap water. The mice received humane care, and experiments were carried out according to the criteria outlined in the Guide for the Care and Use of Laboratory Animals and with the approval of the Animal Care and Use Committee of Hebei Medical College.

The mice were assigned to 6 groups (n=6 per group). We infected 2 groups of mice with PBS (20 μl), 2 groups with pAAV-G-CMV-SV40-NC-RFP (20 μl containing approximately 1×10^11^ AAV virions, produced by Applied Biological Materials) and 2 groups with pAAV-G-CMV-SV40-CCR1-RFP (20 μl containing approximately 1×10^11^ AAV virions, produced by Applied Biological Material) for 18 days. Then, one randomly chosen mouse in each group was sacrificed to verify the CCR1 overexpression. Then, we infected the 3 groups of mice with RSV-GFP (2×10^6^ PFU per mouse) for three days, and three other groups were treated with phosphate-buffered saline (PBS) for 3 days. The expression of the chemokine CCL5 in alveolar lavage solution was detected by ELISAs. Lung RNA was extracted, and the expression of the RSV-N and F genes was detected by qRT–PCR. Total lung imaging (IVIS Spectrum CT, PerkinElmer) was performed to observe the pulmonary RSV load. Hematoxylin and eosin (H&E) staining was performed to detect pulmonary inflammation.

### Cells and virus

Vero cells were grown in Dulbecco’s modified Eagle’s medium supplemented with 10% fetal bovine serum (FBS) and antibiotics (penicillin and streptomycin, Solarbio, China) in a humidified 5% CO_2_ atmosphere at 37°C. Hep-2 cells and A549 cells were grown in RPMI 1640 medium supplemented with 10% FBS and antibiotics in a humidified 5% CO_2_ atmosphere at 37°C. The RSV-GFP A strain, which was kindly provided by Professor He Jinsheng (Beijing Jiaotong University), was amplified in Vero cells.

### Viral infection and virus titer assays

Cells were washed once with PBS and incubated with RSV-GFP at an MOI=1 or mock-infected (with serum-free RPMI 1640 medium alone) for 2 hours. The supernatant was aspirated after viral adsorption, and the cells were maintained in RPMI 1640 supplemented with 2% (v/v) FBS for the indicated times. RSV-GFP was diluted tenfold and used to infect Hep2 cells as described above. Three days later, the titers were measured by plaque-forming assay, and the results are expressed as PFU/milliliter cell lysate or by the number of GFP signals.

### RNA extraction and qRT–PCR

Total RNA was isolated with RNAiso Plus reagent (9101; TaKaRa, Japan). First-strand cDNA was generated with a PrimeScript RT reagent kit (047A; TaKaRa, Japan). Real-time PCR was performed with an ABI Prism 7500 sequence detection system (Applied Biosystems, USA) using Power Up SYBR Green Master Mix (A25742; Applied Biosystems, USA). ACTB was employed as the endogenous control for mRNA relative expression.

### Plasmids and siRNAs

CCL5 gene sequences were synthesized and subcloned into a pCAGGS-HA plasmid, and CCR1, CCR3 and CCR5 gene sequences were synthesized and cloned into a pCDNA3.1-FLAG, pCDNA3.1-HA and pCDNA3.1-HIS plasmid, respectively, by Simon Spirit Biological Technology Company (China). Gαi and β-arrestin 2 gene sequences were synthesized and subcloned into CV155-FLAG and GV143-FLAG vectors (GENE, China), respectively. All the plasmids were extracted with a QIAGEN Plasmid Amp Kit Mini and split after the concentration of each was measured to determine the dose to use in subsequent experiments. The sequences of siCCR1, siCCR5 and siGαi were synthesized by the Gemma Gene Company, and si-β-Arrestin2 was synthesized by Sangon Biotech (China). The following sequences were used:

siCCR5, sense, 5’GCCCAGUGGGACUUUGGAATT3’, and

antisense, 5’UUCCAAAGUCCCACUGGGCTT3’;

siGαi #1, sense, 5’GCUGCUGAAGAAGGCUUUATT3’, and

antisense, 5’UAAAGCCUUCUUCAGCAGCTT3’;

siGαi #2, sense, 5’GCGGAAGAAGUGGAUUCAUTT3’, and

antisense, 5’AUGAAUCCACUUCUUCCGCTT3’;

siITPR3 #1, sense, 5’CCUGCACAUGAAGAGCAACAATT3’, and

antisense, 5’UUGUUGCUCUUCAUGUGCAGGTT3’; and

siITPR3 #2, sense, 5’CGUGUUUAUCAACAUCAUCAUTT3’, and

antisense, 5’AUGAUGAUGUUGAUAAACACGTT3’.

### Cell transfection

Before transfection, A549 **c**ells were seeded in six-well plates. Upon reaching 70-80% confluence, the A549 cells were transiently transfected with the corresponding plasmids or siRNAs by Lipofectamine2000 (Invitrogen, USA). After 48 hours, the efficiency of each transfection was determined by Western blotting.

### Immunofluorescence

A549 cells were seeded on coverslips and transiently transfected with the corresponding plasmids with Lipofectamine 2000 (Invitrogen, USA). Twenty-four hours later, the cells were washed in PBS, fixed in 4% formaldehyde for 15 min, and permeabilized with PBST (0.5% Triton X-100 in PBS) for 10 min at room temperature. The cells were washed three times with PBS and incubated in 10% normal goat serum (Solarbio, China) for 30 min. Primary antibodies (anti-CCR1, anti-CX3CR1) (PA1-41062, Thermo, USA; 13885-1-AP, Proteintech, China) were diluted with PBS containing 10% goat serum, and the cells were incubated with primary antibodies for 1 hour at room temperature. After three washes, the cells were incubated with secondary antibodies containing Alexa 488-labeled goat anti-rabbit (ab150080, 1:500, UK). Coverslips were mounted on the glass slides with DAPI Fluoromount-G (Southern Biotech, USA) for fluorescence microscopy or confocal imaging (Leica TCS SP5).

### RNA-seq analysis

Total RNA from A549 cells was isolated and employed for RNA-seq analysis. Construction of the cDNA library and sequencing were performed by Shanghai Biotechnology Corporation using an Illumina HiSeq 2000 system. High-quality reads were mapped with *Homo sapiens* genome hg38 using HISAT2 version 2.0.4. The expression level of each gene was standardized to fragments per kilobase of exon model per million mapped reads (FPKM) using StringTie version 1.3.0 and the trimmed mean of M values (TMM) (P<0.05, fold change, ≥2).

### Western blot analysis

Total cell lysates were prepared with a radioimmunoprecipitation assay (RIPA) in an ice bath for 30 min and supplemented with 1 mM phenylmethylsulfonyl fluoride (PMSF; BL507A; BioSharp, China) and phosphate inhibitors (P1260; Solarbio, China). The protein concentration was determined with a NanoDrop 2000c spectrophotometer (EW-83061-12; Thermo Scientific, USA). A total of 20-30 μg of protein was separated by 12% sodium dodecyl sulfate–polyacrylamide gel electrophoresis (SDS–PAGE), and the bands were transferred to a polyvinylidene fluoride (PVDF) membrane (IPVH00010; Millipore, USA), which was blocked with 5% nonfat milk. After incubation with antibodies specific to RSV-M2-1 (ab94805, Abcam, UK), GFP (ABclonal, China), β-actin (AC026, ABclonal, China), ATP1A1 (14418-1-AP, Proteintech, China), HSP70 (10995-1-AP, Proteintech, China), HSP90 (13171-1-AP, Proteintech, China), CDC2 (19532-1-AP, Proteintech, China), CCR1 (DF2710, Affinity, China), CCR3 (DF10205, Affinity, China), CCR5 (69n9918, Affinity, China), GAPDH (5174S, CST, USA), p-ERK (76299, Abcam, UK), ERK (184699, Abcam, UK), p-STAT3 (CY6566, Abcam, China), or STAT3 (CY5165, Abcam, China), the blots were incubated with horseradish peroxidase (HRP)-labeled goat anti-mouse IgG (Abgent, USA) or HRP-labeled goat anti-rabbit IgG (Abgent, USA) and then detected with Western Lightning plus-ECL reagent. GAPDH or β-actin was used as a loading control for immunoblotting.

### Flow cytometry

A549 cells were transfected with 500 ng of plasmid containing the target gene CCR1 and infected with RSV for 24 hours, and the control group was set up at the same time. All supernatant and adherent cells were collected in a centrifuge tube and centrifuged at 1000 rpm for 5 min. Then, the supernatant was removed and washed in PBS twice. The cell concentration was diluted with 1× binding buffer. A total of 1×10^6^ cell suspension samples (approximately 100 μL) were placed in a flow cytometer in tubes with 5 μL of PE-annexin V (559763, BD, USA) and 5 μL of 7-AAD (559925, BD, USA) for staining, and the cells were incubated with the dyes at 25°C for 15 min in the dark. After adding 1× binding buffer (400 μL) to each tube, the suspension sample were mixed and measured by flow cytometry.

### IP

IP assays were performed according to the instructions of a Pierce classic magnetic IP/co-IP kit. Briefly, the cells were lysed in specific buffer on ice for 30 min. The supernatant protein was then incubated overnight with the corresponding antibody on a rotator at 4 °C. The next day, Pierce protein A/G beads were washed with specific buffer three times. The supernatant-antibody mixture and the beads were then coincubated on a rotator at room temperature for 2 hours, washed with lysis buffer and PBS, and boiled for 10 min. The samples were subjected to SDS–PAGE and Western blot analysis, and the target proteins were detected using the corresponding antibodies.

### Virus entry and adhesion

RSV was added to CCR1-overexpressing A549 cells, incubated at 4 °C for 1 hour to allow the RSV to adsorb onto cells, and then cultured at 37 °C for 30 min to initiate virus entry into cells. Total RNA in the cells was immediately extracted, and the expression of the RSV-N and F genes was detected by qRT–PCR.

### Statistical analysis

Data from three independent experiments are expressed as the mean ± standard deviation (SD). GraphPad Prism software was used for statistical analyses.

## DATA AVAILABILITY

The RNA-seq data set is available in the Gene Expression Omnibus (GEO) and can be accessed via the GSE series number GSE145371.

## ACKNOWLEDGMENTS

Thanks to Professor He Jinsheng of Beijing Jiaotong University for providing RSV with GFP label. The work was supported by grants from the National Natural Science Foundation of China (81971474 and 82071787) and Key Project of Natural Science Foundation of Hebei Province (C2021206011).

## AUTHOR CONTRIBUTIONS

All authors have read and approved the article. W.L., M.CQ., L.J. and L.M. conceived and supervised the study. W.L., M.CQ. and L.J. wrote the paper. X.L., M.LT., W.XL. and G.X. collected samples and data. L.J., X.L. and M.LT. performed the experiments. L.M., W.JC., W.XL. and G.X. contributed to the design of the study and data analyses.

---

**Supplementary Fig.**
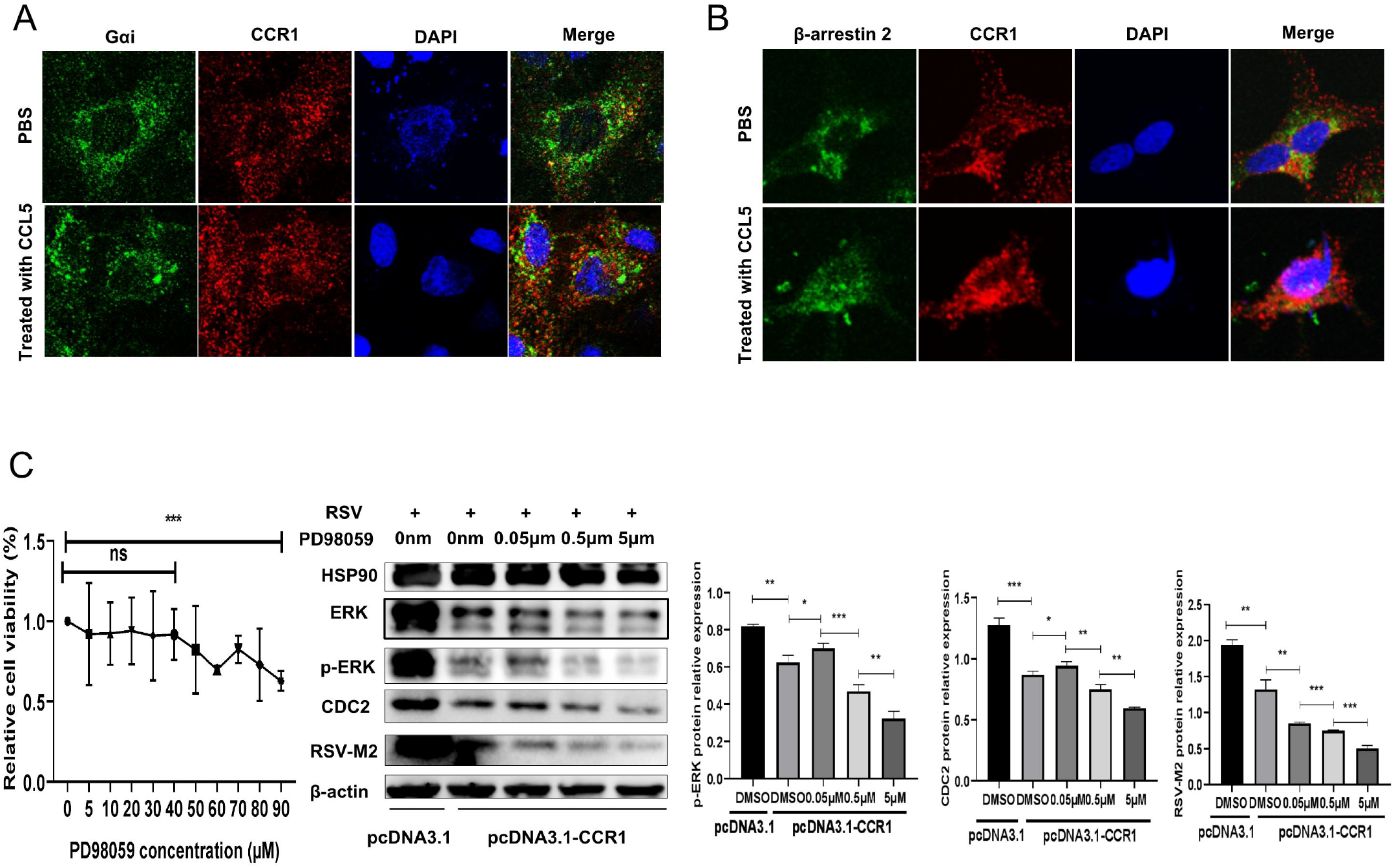
(A, B) Following overexpression of CCR1 and Gαi or β-arrestin2 in A549 cells, the recruitment of Gαi or β-arrestin2 by CCR1 was observed by confocal microscopy. (C) CCR1-overexpressing A549 cells were pretreated with different concentrations of an ERK inhibitor (PD98059) and then infected with RSV. The protein expression levels of CDC2, p-ERK and RSV-M2 were detected by Western blotting. (Unpaired t test, \**P*<0.05, \*\**P*<0.01, \*\*\**P*<0.005, and \*\*\*\**P*<0.001)

